# Unraveling the inbreeding depression patterns in a self-pollinated eucalyptus population

**DOI:** 10.1101/2025.02.03.636226

**Authors:** Filipe Manoel Ferreira, João Amaro Ferreira Vieira Netto, Guilherme Ferreira Melchert, Thiago Romanos Benatti, José Wilacildo de Matos, Fabiana Rezende Muniz, Itaraju Junior Baracuhy Brum, André Vieira do Nascimento, Leonardo Lopes Bhering, Kaio Olimpio da Graças Dias, Evandro Vagner Tambarussi

## Abstract

Genetic improvement greatly contributed to the success of the forest industry in Brazil. While past selection efforts in *Eucalyptus* spp. have yielded satisfactory genetic gains, the response to selection in the last decade has fallen below expectations. Recent research suggests that inbreeding-based selection strategies, well-established in crops such as rice and maize, could be adapted to enhance perennial species such as guava and can be expanded to forest species, such as eucalypt. In this context, 20 elite *Eucalyptus* spp. genotypes were self-pollinated, producing 30 progenies per family. A total of 600 individuals were planted in an experimental trial and evaluated for growth traits at 3 years of age. Both the progeny and the 20 parent genotypes were genotyped using SNP chips. This research aims to unravel the patterns of inbreeding depression in this *S*_0:1_ population of *Eucalyptus* spp., by examining autozygosity and genomic inbreeding patterns, estimating inbreeding depression (*ID*) and genetic parameters, and studying the impact of the unbalance between selfed and crossed individuals on the inbreeding depression estimator. The results revealed that dominance variance accounted for a notable portion of the phenotypic variation, demonstrating the significance of non-additive genetic effects in diameter at breast height (*DBH*). *ID* was evident, with reductions in *DBH* observed in most families as homozygosity increased. Simulations highlighted that unbalanced sample sizes of selfed and crossbred individuals could bias estimates of *ID*. By integrating genomic data and advanced quantitative methods, this study brings new information into the genetic consequences of self-pollination in eucalyptus, offering a foundation for managing inbreeding and enhancing genetic gains in perennial breeding.

## 1. Introduction

The use of inbred lines for single-cross hybrid production was one of the most significant and impactful breeding techniques for outcrossing species, dramatically enhancing genetic gains in maize breeding programs (Hallauer et al., 2010; Bernardo, 2021). Despite the success of this technique, forest companies were slow to adopt this approach to produce superior hybrids. Few forest breeders have investigated the effects of inbreeding in partially inbred lines of *Eucalyptus* spp. or proposed methods for their commercial application in breeding programs (Ramalho et al., 2023). This delay arises because of the long time required to achieve high endogamy, and the lack of knowledge about the heterozygosity and inbreeding depression already present in the breeding populations, as well as the potential effects of dominance (heterosis) from crosses between partially inbred plants. Studying genomic patterns of homozigosity in inbred populations and understanding inbreeding depression and heterosis processes have the potential to increase the rate of genetic gain (Jighly et al., 2019).

Traditional tree breeding programs typically involve crossing high-performing heterozygous genotypes (i.e. clones) to produce highly productive and complementary progenies. This approach enables breeders to select the best hybrids from the best progenies and fix these genotypes via clonal propagation (Resende and Barbosa, 2005). However, the correlation between genotypes selected in progeny trials and their corresponding clones in clonal trials tends to be low (Lima et al., 2011), thereby limiting the ability to identify and propagate highly productive clones. This challenge arises from several factors: poor representation of crosses due to the small number of individuals evaluated in progeny field trials and limited observations (Resende et al., 2017b), genetic competition (Ferreira et al., 2024, 2023), the absence of suitable mating designs to capture both additive and non-additive effects effectively (Vitezica et al., 2013), the lack of information about the mating system (Grattapaglia et al., 2012), and the disregarded inbreeding within the population (Charlesworth and Willis, 2009).

Inbreeding occurs when genetically related individuals mate, increasing homozygosity and reducing genetic diversity. In populations with inbred crosses, genetic relatedness between individuals can be higher than expected. For example, assuming half-sib relationships based on pedigree matrix in open-pollinated trials can result in biased genetic estimates, as the actual degree of kinship is often higher than presumed due to inbreeding (Tambarussi et al., 2018). Consequently, genetic variance within the population is underestimated, leading to flawed assessments of heritability and breeding values. Furthermore, some species have a mixed matting system (Goodwillie et al., 2005; Duminil et al., 2009; Winn et al., 2011) such as many of the genus *Eucalyptus* (Tambarussi et al., 2018; Griffin et al., 2019; Yeh et al., 2011).

The one-locus model with two alleles (M/m) can help to elucidate the genetic basis of inbreeding depression: by assuming that the frequencies for homozygote genotypes MM and mm are D and R, respectively, and the frequency for heterozygote genotype Mm is H, we can expect that the mean population fitness for an initial population without selfing is *ω*_0_ = *Dv*1 + *Hv*2 + *Rv*3, being *v*1, *v*2, and *v*3 the fitness value of the possible genotypes. After one generation of selfing we expect that the mean population fitness is 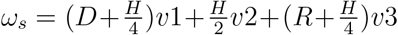 due to recombination. Therefore, inbreeding depression occurs if *ω*_0_ *> ω*_*s*_, which depends on the value of the heterozygote relative to the two homozygotes and the dominance of one allele over the other, which can be represented by the higher fitness of the heterozygote related to the mean of homozygotes 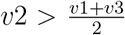 (Cheptou, 2024). In a multilocus, after one generation of selfing, it is expected that at least 50% of the alleles will be homozygous due to inbreeding *F*_*s*_ = 100[0.5(1 + *F*_0_)], being *F*_0_ the pre-existing inbreeding. Thus, inbreeding depression refers to the reduction of biological fitness in a population due to crossing of related individuals or selfing (Charlesworth and Willis, 2009).

Cockerham and Weir (1984) proposed a genetic model to obtain the total genetic variance as a function of the inbreeding coefficient for mixed matting species. However, it requires an elaborated cross design and the evaluation of numerous family and parental structures to capture the covariances of all parent and first-generation relatives, which are influenced by both cross-and self-fertilization. Currently, genotyping using Single Nucleotide Polymorphisms (SNP) arrays has become increasingly affordable being a powerful tool to help partition genotypic variance and predict genotypic values in populations under inbreeding (Silva-Junior et al., 2015). This allows for more comprehensive analyses that incorporate portions of the genome that can be in linkage disequilibrium with the phenotypic trait (Grattapaglia, 2017). Based on that, we used molecular data from a *S*_0:1_ population of *Eucalyptus* spp. to: i) investigate genome-wide autozygosity patterns and genomic inbreeding level based on homozygosity-by-descent (HBD) segments; ii) estimate genetic parameters for a self-fertilized population of *Eucalyptus* spp.; iii) estimate inbreeding depression levels in the family level; and iv) evaluate the impact of unbalance among the number of self-fertilized and cross-fertilized individuals per family on the estimates of inbreeding depression using a simulated dataset that emulates our real dataset.

## 2. Material and Methods

### 2.1. Plant material

A total of 20 elite *Eucalyptus* spp. genotypes, here called families, including 11 derived from crosses between *Eucalyptus urophylla* and *Eucalyptus grandis* (“urograndis”), 5 from *Eucalyptus urophylla*, 1 from *Eucalyptus grandis*, and 3 from *Eucalyptus* spp., were subjected to self-pollination. After germination, 30 individuals were collected from each family, resulting in 600 individuals (progenies). These were established in a field experiment using a randomized complete block design, with one plant per plot and 30 blocks. Additionally, 8 parent trees and 5 commercial clones were included as checks. The 600 progenies and the 20 parents were genotyped using Illumina Infinium 60k BeadChips panels Silva-Junior et al. (2015), and phenotyped for Diameter at Breast height (DBH, cm) at 3 years of age.

### 2.2. Quality Control

To ensure the integrity of genotype data for subsequent analyses, we implemented a quality control protocol using `PLINK v1.9` (Purcell et al., 2007). SNPs with a call rate below 90% were excluded using the `–geno 0.1` parameter. Also, SNPs with a minor allele frequency (MAF) less than 0.05 were removed by applying the `--maf 0.05` parameter.

### 2.3. Paternity test

Molecular markers (SNP) were coded as 0 and 2 for homozygotes and 1 for heterozygotes. High polymorphic markers were selected to conduct the paternity test on the software Cervus (Marshall et al., 1998; Kalinowski et al., 2007). Cervus employs the maximum likelihood method to estimate allele frequencies, calculate likelihoods, and assign parentage based on the LOD score, which compares the probability of a true parent with that of a random candidate. Additionally, it measures the difference between the LOD of the most likely parent and the second candidate, increasing confidence in the assignment. Monte Carlo simulations were conducted to validate the assignments, considering a 95% significance level and minimizing potential genotyping errors. After the paternity test, we could identify individuals resulted from cross-and self-fertilization.

Despite controlled self-fertilization, cross-fertilized individuals were observed for some families. This allowed us better understand how inbreeding influences phenotypic performance by estimating the *ID*_%_ at the family level. For this, we defined a family as a set of individuals that have the same parental male, since there is a greater representation of the male parent in relation to the female one. Therefore, we have a total of 26 families composed by different proportions of the sections Exsertaria, Maidenaria, and Transversaria (Supplementary Material - Figure S1). Of them, we have 16 families that simultaneously have selfed and crossed individuals evaluated on the trial (Figure 2).

### 2.4. Estimating genetic parameters

We employed an additive-dominant genomic best linear unbiased prediction (AD-GBLUP) linear mixed model to estimate genetic parameters and predict genetic effects. The linear mixed model can be represented as:

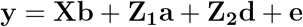

where **y** represents the (*n×*1) vector of phenotypic observations; **X** is the (*n× r*) design matrix linking fixed effects to each individual, where *r* is the number of fixed effects (blocks); **b** is the (*r ×* 1) vector of fixed effects; **Z**_**1**_ is the (*n × q*_*a*_) design matrix associating phenotypic records to additive genetic effects, where *q*_*a*_ is the number of individuals in the population; **a** denotes the (*q*_*a*_ *×*1) vector of additive genetic effects for individuals 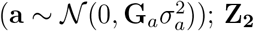 is the (*n × q*_*d*_) design matrix associating phenotypic records to dominance genetic effects, being *q*_*d*_ the number of individuals with dominance effects modeled; **d** denotes the (*q*_*d*_ *×* 1) vector of dominance genetic effects for individuals 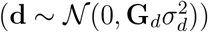; and **e** is a (*n×*1) vector of residual effects 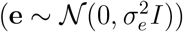. Furthermore, **G**_*a*_ is the (*q*_*a*_ *× q*_*a*_) genomic relationship matrix for additive effects, **G**_*d*_ is the (*q*_*d*_ *× q*_*d*_) genomic dominance relationship matrix, *I*_*n*_ is the identity matrix of order *n*, 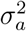 is the additive genetic variance, 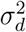 is the dominance genetic variance, and 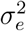 is the residual variance.

The genomic relationship matrix (**G**) was calculated as follows VanRaden (2008) :

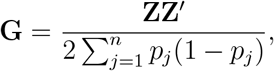

where **Z** is a matrix of centered marker genotypes, defined as *Z*_*ij*_ = {2 − 2*p*_*j*_, 1 − 2*p*_*j*_, −2*p*_*j*_} Vitezica et al. (2013). Here, *Z*_*ij*_ is the element of the *i*^*th*^ row and *j*^*th*^ column of the marker matrix, and *p*_*j*_ represents the allele frequency of the second most frequent allele at locus *j*. The construction of **Z** involves transforming the nucleotide marker genotypes into values of 0, 1, and 2, where 0 corresponds to the most frequent homozygote, 1 to the heterozygote, and 2 to the second most frequent homozygote.

The dominance genomic relationship matrix (D) was obtained following Vitezica et al. (2013):

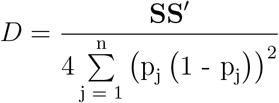

where **S** is a dominance marker matrix, defined as 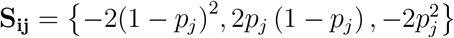, being *S*_*ij*_ the elements of the ith row and jth column of the *S* matrix and *p*_*j*_ the allele frequency of the jth marker.

We used the components of variance extracted from the AD-GBLUP to estimate the narrow sense heritability 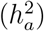, coefficient of determination for dominance effects (*d*^2^), the broad sense heritability (*H*^2^), and the mean selection accuracy for additive (*r*_*âa*_) and dominance 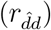 effects, defined as follows:

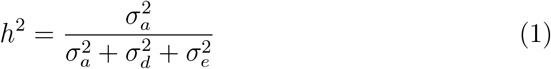

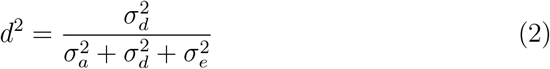

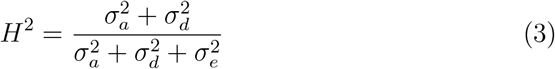

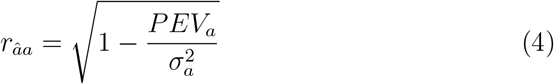

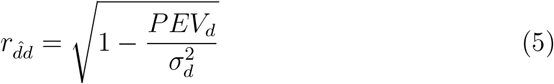

where *PEV*_*a*_ and *PEV*_*d*_ denote the prediction error variance derived from the diagonal of the inverted coefficient matrix for the *a* and *d* effects, respectively. The expected response to selection, in percentage, *R*%, by selecting the best ranked 30 genotypes based on the genotypic value and the 3 most endogamic individuals of the 10 most endogamic families is given by:

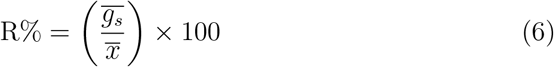

where 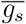 is the mean genotypic value of the selected genotypes, and 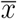 is the mean phenotypic value.

### 2.5. Estimation of inbreeding coefficients

We estimated the inbreeding coefficients of each self-fecundated individual based on the sequence of consecutive homozygous-by-descendent segments (HBD). Alemu et al. (2021) compared several methods for measuring inbreeding in cattle and concluded that HBD makes better use of the information from neighboring markers. HBD segments were detected with the RZooRoH v.0.3.1 package in R (Bertrand et al., 2019). The RZooRoH package applies a hidden Markov model to detect HBD segments, alongside independent segments following Druet and Gautier (2017). The frequency and length of HBD segments serve as indicators of historical inbreeding. Longer segments imply more recent inbreeding, while shorter ones point to older inbreeding events.

A model was configured with the number of HBD classes equal to *K* = 10, with rate *R*_*k*_ equal to 2, 4, 8, 16, 32, 64, 128, 256, and 512 generations. Segments longer than 512 generations were classified as non-HBD. The lengths of HBD segments follow an exponential distribution, where *R*_*k*_ determines the expected length of HBD segments (Thompson, 2013). Each *R*_*k*_ value approximates twice the number of generations since the inbreeding event occurs. Therefore, they represent ancestry from 1, 2, 4, 8, 16, 32, 64, 128, or 256 generations ago. The realized inbreeding coefficient based on the HBD segments *F*_*HBD*_ was estimated as the probability of belonging to any HBD classes averaged over the whole genome (Bertrand et al., 2019; Druet and Gautier, 2017).

### 2.6. Inbreeding depression

In the crosses, there is greater representation of the male parent in relation to the female one. Therefore, we modeled the linear relationship between the *F*_*HBD*_ and the DAP for each male parent. For the male parent *i*, the model is given by:

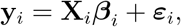

where **y**_*i*_ is the vector of DAP values for the *n*_*i*_ genotypes in family *i*; **X**_*i*_ is the design matrix containing a column of ones for the intercept and acolumn of endogamy coefficients (*F*_*HBD*_) for the genotypes; ***β*** _*i*_ = [*β*_0*i*_, *β*_1*i*_]^*T*^is the transposed vector of regression coefficients, with *β*_0*i*_ representing theintercept and *β*_1*i*_ the slope (effect of *F*_*HBD*_ on DAP);***ε***_*i*_ is the error vector, assumed to follow a normal distribution, ***ε***_*i*_ ∼ 𝒩 (**0**, *σ*^2^**I**). The coefficient of determination (*R*^2^) was computed for each regression.

Based on the mean *F*_*HBD*_, we selected the 10 most endogamic families and the 3 most endogamic individuals within these families, totaling 30 high endogamic individuals.

Inbreeding depression on DBH, in percentage (*ID*(%)), was estimated for families that included at least one inbred individual considering the derivations of Cockehan (1986) later presented by Resende (2002) and applied by Simiqueli et al. (2018) for mixed matting species:

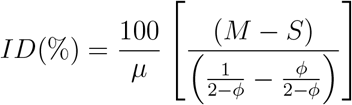

where *µ* is the average of *DBH, X*_*C*_ is the phenotypic mean for individuals from cross-fertilization, considering *F*_*HBD*_ = 1*/*(2 − *ϕ*) and *X*_*S*_ is the phenotypic mean for individuals from self-pollination, *ϕ* is the family self- fertilization coefficient in populations with mix mating system.

### 2.7. Simulations

Our data were simulated to emulate the original data. After testing many values for the simulation parameters (Gaynor et al., 2021), we achieve an overlay of 89% between the real and simulated data (Supplementary Material - Figure S2).

To investigate how the imbalance between the individuals derived from self-fertilization and those from crossbreeding impacts the accuracy of the inbreeding depression estimator (*ID*), a simulation was carried out to test the estimator’s performance under different imbalance scenarios. The goal was to determine how robust the estimator is when faced with varying proportions of selfed and crossed individuals. The simulation began by generating two founder populations of a hypothetical species. These populations underwent recurrent selection, after which selected individuals from each population were crossed to create a hybrid population. This hybrid population, composed of improved individuals from both founders, was further subjected to recurrent selection. In the final generation, individuals were either self-fertilized or crossbred (Figure 1), replicating a process similar to what was observed in the real dataset used in this study.

**Figure 1:**
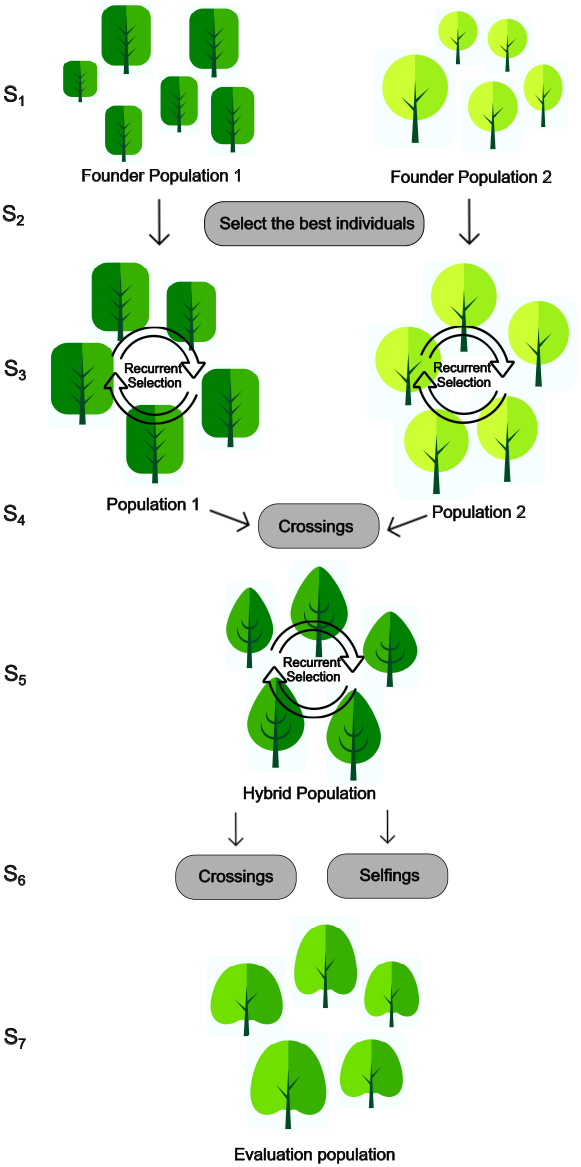
Simulation workflow for emulating a *S*_0:1_ *Eucalypts* spp. breeding population. A total of seven steps (S) were applied. *S*_1_: Two founder populations were generated, each belonging to a different species, these populations underwent random mating for 100 cycles, without any selection. *S*_2_: The top 10 individuals from each founder population were selected. *S*_3_: The individuals selected in the previous step were subjected to one cycle of recurrent selection separately for each population. *S*_4_: Crosses were performed between the top 20 individuals of the first population and the top 20 of the second population, resulting in a hybrid population. *S*_5_: The hybrid population underwent 10 cycles of recurrent selection. *S*_6_: The best individual from *S*_5_ population was selfed and crossed. *S*_7_: From this process, 1500 (15 combinations x 100 repetitions) populations were created with combinations of selfed in relation to crossed individuals ranging from 29/1 to 15/15. Based on this workflow, we achieved 89% overlap between real and simulated data.

To evaluate the effect of imbalance, several families were simulated with different proportions of selfed and crossed individuals. These combinations ranged from 15 selfed and 15 crossed individuals (15/15) to 29 selfed and 1 crossed individual (29/1), resulting in 15 unique scenarios. For each combination, *ID* was estimated, and this process was repeated 100 times to capture the mean, variance, minimum, and maximum values for each scenario. The root mean square error (*RMSE*) was calculated to measure how much the simulated *ID* values (*ID*_*s*_) deviates from the *ID* of the most balanced scenario based on the genotypic values of the real data (*ID*_*g*_). The *RMSE* is computed as:

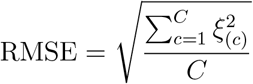

where *ξ*_(*c*)_ is the esimatated bias or error for the *ID* comparing the *ξ*_(*c*)_ = *ID*_*s*_ − *ID*_*g*_, and *C* the number of cycles of simulation in stage S7 (100, in our case) (Figure 1). The relative bias in percentage 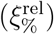 is computed as:

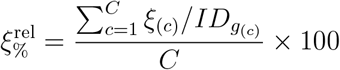

Due to the stochastic nature of the simulation, in general it is not possible to generate a population with genetic parameters that perfectly match those of the real population. Therefore, the simulation was run 5,000 times, and the population with the highest overlap between the phenotypic distribution of the simulated and real populations was selected for this study. The genetic parameters of the simulated populations were set to reflect the traits of the real population analyzed in this study (Table 2).

**Table 1:** Number of individuals from self- and cross-fertilization for the male parent.

| Male parent | Selfed | Crossed |
| --- | --- | --- |
| G2* | 12 | 5 |
| G11 | 16 | 5 |
| G10 | 7 | 3 |
| G1 | 27 | 27 |
| G24 | 32 | 4 |
| G7 | 17 | 30 |
| G22 | 27 | 9 |
| G21 | 10 | 1 |
| G29 | 4 | 1 |
| G23 | 29 | 9 |
| G20 | 12 | 15 |
| G5 | 26 | 2 |
| G4 | 17 | 5 |
| G25 | 8 | 5 |
| G8 | 18 | 6 |
| G27 | 9 | 5 |

\* Parents' names were randomly generated ranging from 1 to 30.

**Table 2:** Variance components and genetic parameters estimated via the AD-GBLUP model.

| Parameter | Estimate | Standard Error |
| --- | --- | --- |
| $\sigma_a^2$ | 4.98 | 1.86 |
| $\sigma_d^2$ | 2.86 | 1.30 |
| $\sigma_e^2$ | 12.62 | 0.84 |
| $\sigma_p^2$ | 20.46 | |
| $h_a^2$ | 0.24 | 0.08 |
| $d^2$ | 0.14 | 0.06 |
| $H^2$ | 0.38 | 0.06 |
| $r_{\hat{a}a}$ | 0.59 | |
| $r_{\hat{d}d}$ | 0.43 | |
| $R_g(\%)$ | 6.99 | |
| $\overline{X}_s$ | 15.70 | |
| $\overline{X}_0$ | 12.65 | |

where $\sigma_a^2$ denotes the additive genetic variance, $\sigma_d^2$ denotes the dominance genetic variance, $\sigma_e^2$ denotes the residual variance; $\sigma_p^2$ is the phenotypic variance; $h_a^2$ is the narrow sense heritability; $d^2$ is the coefficient of determination for dominance; $H^2$ denotes the broad sense heritability; $r_{\hat{a}a}$ is the accuracy for additive genetic effects; $r_{\hat{d}d}$ is the accuracy for dominance genetic effects; $R_g$ is the response to selection by selecting the 30 best-ranked individuals via genotypic values ( $a+d$ ).

We performed all analyses in the `R` software environment (Version 4.3.2, R Core Team, 2023). We estimated the inbreeding patterns using the `RZooRoH` package (Bertrand et al., 2019) and we fitted the linear mixed models using the `ASReml-R` package (Version 4.2, The VSNi Team, 2023). We conduct the simulations using the `AlphaSimR` package (Gaynor et al., 2021), and all visualizations using the `ggplot2` package (Wickham, 2016) and the `inkscape` software (Bah, 2011).

## 3. Results

### 3.1. Genetic parameters

According to the Likelihood Ratio Test (LRT), the additive and dominance effects are significant. The additive genetic variance 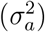 was estimated at 4.98, while the dominance genetic variance 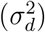 was 2.86. Residual variance 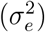 accounted for the largest proportion of the total variance, with an estimate of 12.62 (SE = 0.84). The narrow-sense heritability 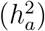 indicates that 24% of the phenotypic variance could be attributed to additive genetic effects. The coefficient of determination for dominance effects (*d*^2^) suggests a contribution of dominance variance to the total phenotypic variance (14%). The broad-sense heritability (*H*^2^), which incorporates both additive and dominance components, was 0.38, evidencing the influence of genetic factors on the DBH (Table 2). The accuracies of the estimated genetic values were 0.59 and 0.43 for additive and dominance effects, respectively. Based on the selection of the top 30 individuals ranked by their genotypic values, *R*_*g*_(%) was 6.99%, which indicates the potential of this self-fertilized population for genetic improvement through selection (Table 2).

### 3.2. Estimation of inbreeding coefficients

We observed a low frequency of individuals presenting *F*_*HBD*_ greater than 0.5 and a high frequency of individuals presenting *F*_*HBD*_ less than 0.15 (Figure(2a). The classes with rate *R*_*k*_ equals 8, 16, and 512 were the most frequent at the genome level (Figure 2b).

**Figure 2:**
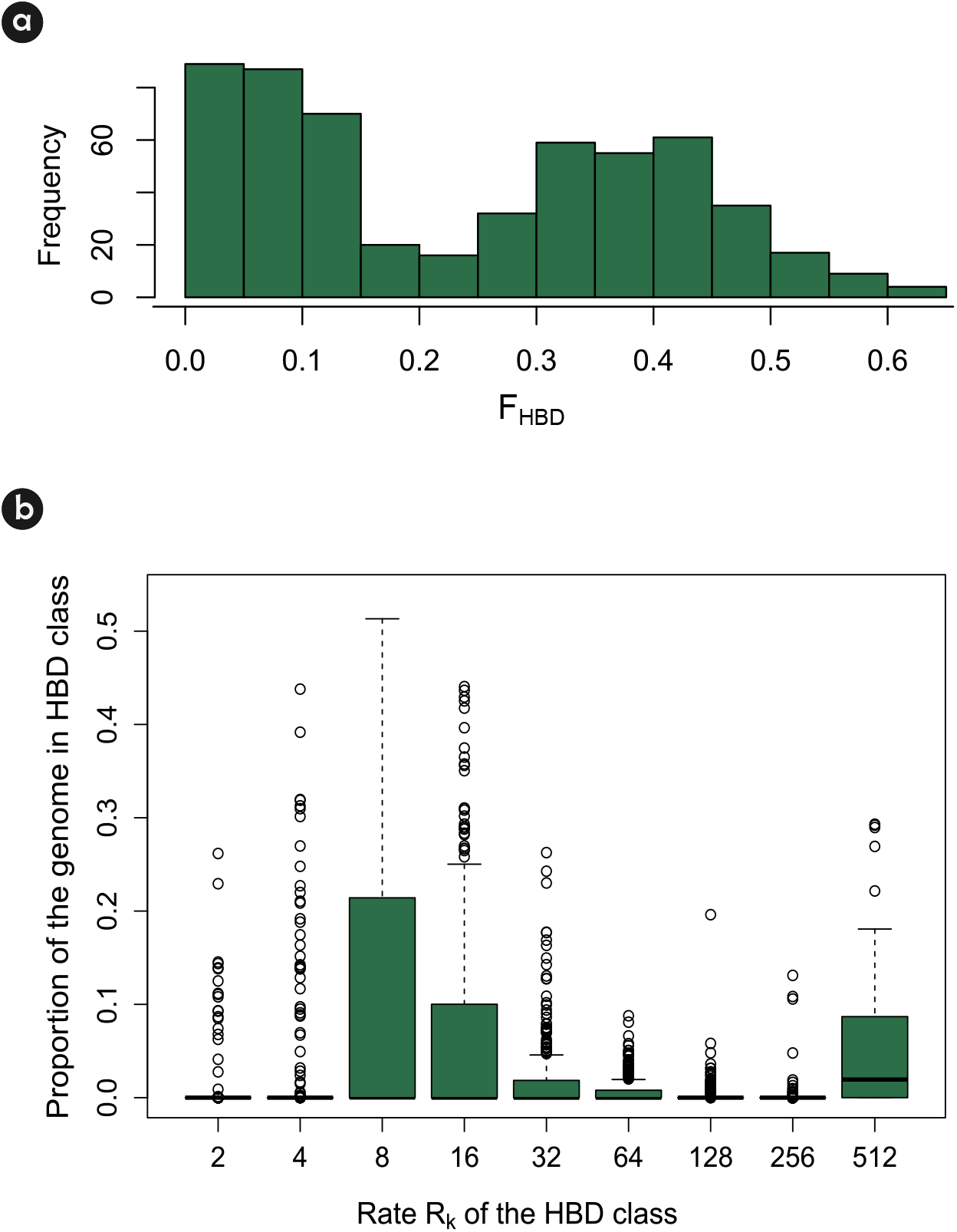
a) Summary statistics of the inbreeding coefficient (calculated as the sum of the contributions of all HBD classes). b) Partitioning of genome-wide autozygosity for a *S*_0:1_ population of *Eucalyptus*. The percentages correspond to individual genome-wide probabilities of belonging to each of the HBD-classes, which are defined by their rate (2, 4, 8, 16, 32, 64, 128, 256, 512).

We estimated the level of autozygosity per HBD class for two different situations: i) based on the 20 most endogamic individuals ranked based on their *F*_*HBD*_ values, and ii) based on the 20 individuals that presented the higher genotypic values (a+d) via the AD-GBLUP model (Figure 3). It is worth mentioning that the class with *R*_*k*_ equals 8 was the most frequent for individuals that presented the highest *F*_*HBD*_ values, while the class with *R*_*k*_ equals 512 was the most frequent for the best performance individuals based on the genotypic value (Figure 3).

**Figure 3:**
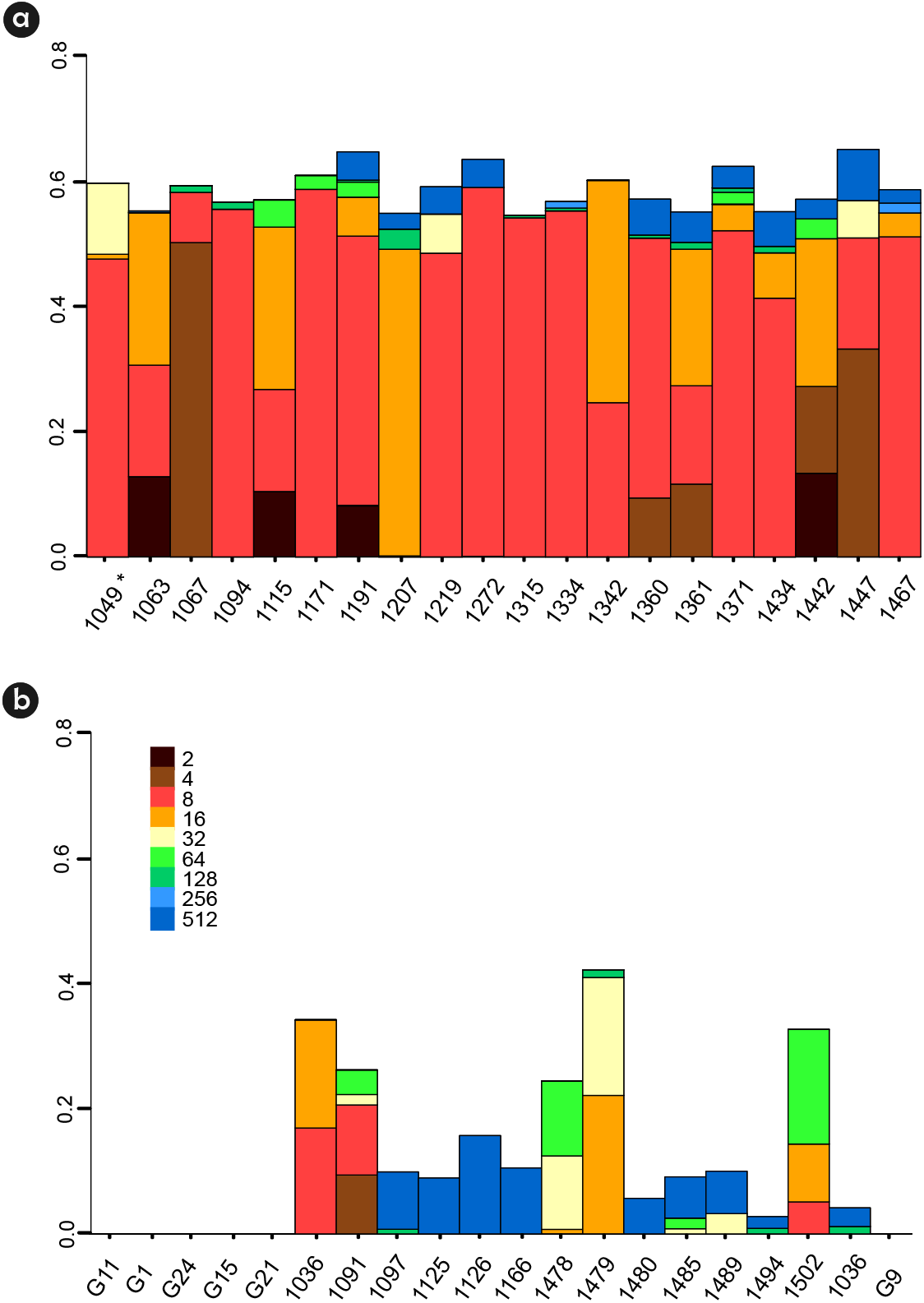
Estimated level of autozygosity per HBD class in the 20 most endogamic individuals based on the *F*_*HBD*_ (a) and on the 20 best-ranked individuals based on the AD-GBLUP model (b). Each colour is associated with a distinct class, defined by its rate (2, 4, 8, 16, 32, 64, 128, 256, 512). The heights of each color bar represent the estimated level of autozygosity associated with the class, and the total height represents the total estimated autozygosity. ^*^ The numbers used for coding the individual’s names were randomly generated from 1000 to 2000 and parent’s names were randomly generated ranging from 1 to 30.

The average *F*_*HBD*_ for all the families was 0.2367. It ranged from 0.02 to 0.35 at the family level. For some families the regression of *F*_*HBD*_ on DAP was able to capture trends, which may indicate inbreeding depression (Supllementary Material - Figure S3).

### 3.3. Inbreeding depression

The average values of the mean of the cross- and self-fertilized populations for DBH were 13.88 and 9.73 cm, respectively. The mean inbreeding depression, considering that the population has a mixed matting system, was 91. 76 %. Inbreeding depression was calculated for families that presented both crossed and selfed individuals. The family G29 showed outbreeding depression, indicating that the selfed individuals presented greater mean than the crossed ones. In all the remaining families, the opposite situation was observed (Figure 4). The *R*^2^ suggests that 42% of the variation in inbreeding depression can be explained by the variation in *F*_*HBD*_.

**Figure 4:**
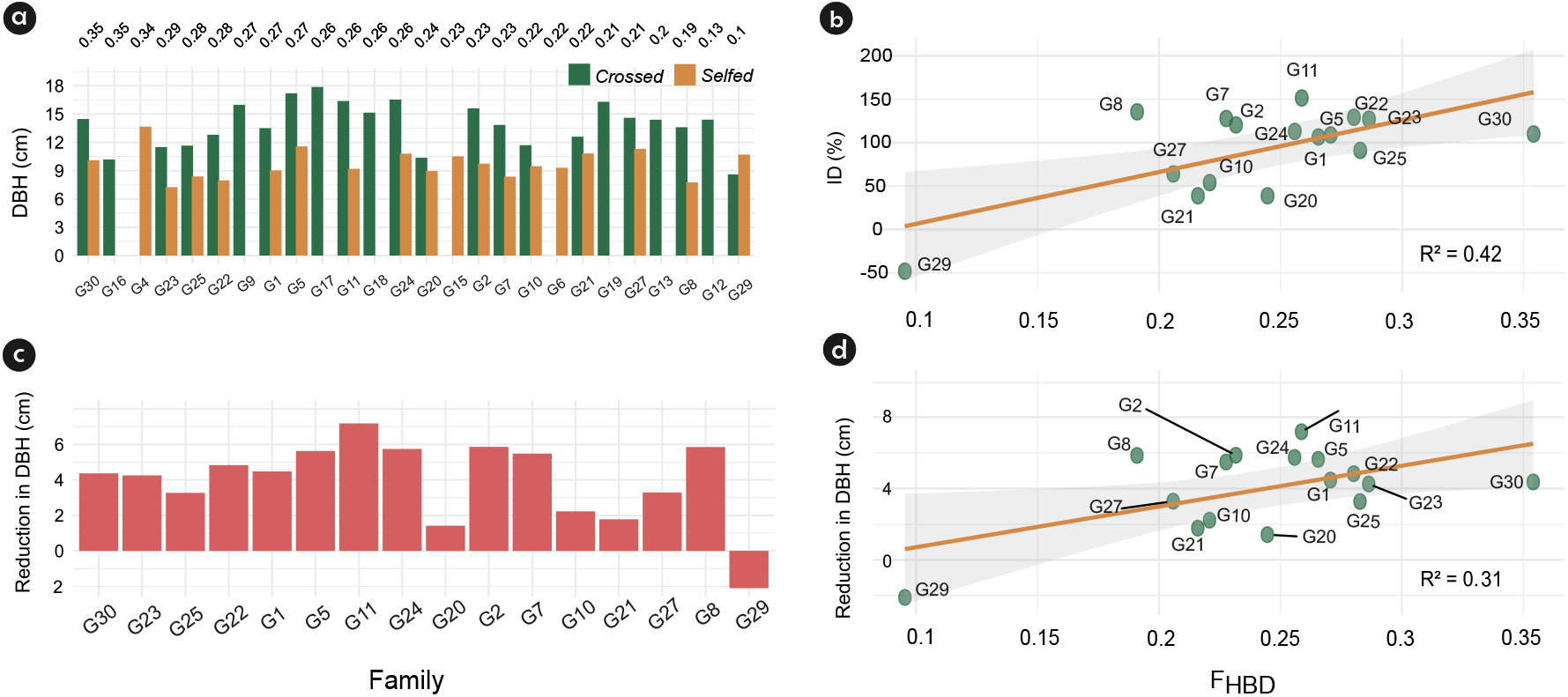
Means and inbreeding depression for a crossed- and selfed-fertilized population. a) Mean of the cross- and self-fertilized population for diameter at breast height (DBH), in cm, for each male parent. b) inbreeding depression in percentage (*ID*(%)), as a function of the inbreeding coefficient obtained by homozygosity-by-descent (*F*_*HBD*_) for families that presented both crossed and selfed individuals. c) Reduction in DBH (cm) due to inbreeding depression (diference among the mean of crossed and selfed individuals). d) Reduction in DBH (cm) as a function of the inbreeding coefficient obtained by homozygosity-by- descent (*F*_*HBD*_) for families that presented both crossed and selfed individuals. *R*^2^ is the coefficient of determination of the regression. The numbers used for coding the parent’s names were randomly generated ranging from 1 to 30.

However, as the proportion of crossbred individuals increases in combinations such as 23.7 through 15.15, the RMSE values drop considerably. This happens because, in more balanced setups, the means for both self-fertilized and crossbred groups are more stable because of larger sample sizes, which reduces variation in the estimates.

### 3.4. Simulation

Figure 5 illustrates the changes in the estimated values of *ID* and variances for different combinations of individuals derived from self- and cross-fertilization. We observed that for all scenarios the *ID* is overestimated (Figure 5a). The most unbalanced combinations of self- and cross-fertilized individuals, such as 29.1, 28.2, and 27.3, exhibit the highest variability (Figure 5a) and Root of the Mean Square Error (*RMSE*) (Figure 5b) in *ID* estimates. In contrast, as the imbalance decreases, for example, 17.13, 16.14, and 15.15, the spread of the estimates narrows to the maximum. As the proportion of crossbred individuals increases reaching a representation greater than 7 (23.7), the RMSE values drop considerably. Therefore, in the evaluated scenarios containing 30 individuals per family, a minimum of 7 crossed individuals should be established among the 30 (23.3%) to ensure a more precise estimate of *ID*.

**Figure 5:**
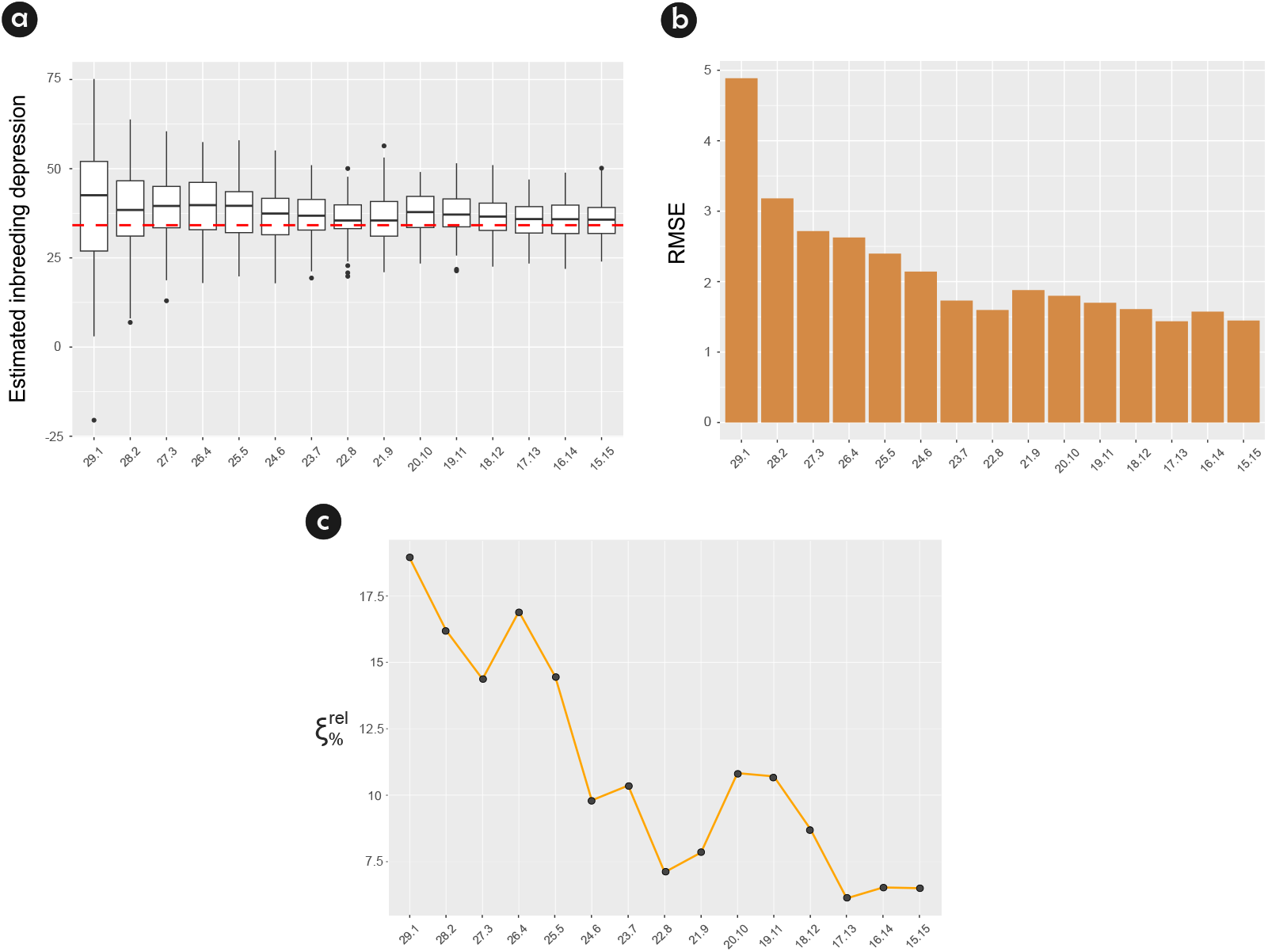
Estimated inbreeding depression (*ID*) and Root Square Mean Error (*RSME*) for different combinations of self- and cross-fertilized individuals based on simulated data. a) On the X-axis, each combination is labeled by the number of self-fertilized individuals followed by the number of cross-fertilized individuals, separated by “.” (that is, ‘29.1’ represents 29 self-fertilized individuals and 1 crossbred individuals). The Y-axis shows the estimated values of *ID*. The dotted red line on the boxplot indicates the true simulated *ID*. b) *RSME* values for the *ID* estimates for different combinations of self- and crossfertilized individuals. c) Relative bias 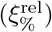 for different combinations of self- and crossfertilized individuals.

We found the mean relative bias of the *ID* estimator equals to 11.02 %. The 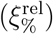 is influenced by the proportion of selfed individuals relative to the crossed ones. When the composition of families is balanced (15.15) 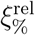 is equal to 6.5% (Figure 5c). As the number of selfed individuals increases, 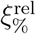 tends to rise, although fluctuations occur in the intermediate ranges (17 to 20 selfed individuals) between 6.12 and 10.8%. A marked increase in 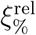 is observed when the number of selfed individuals reaches 23, where 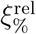 reaches 10.4%. When only one crossed individual remains the 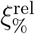 peaks at 18.9% indicating a clear sensitivity to extreme imbalances in family composition (Figure 5c).

## 4. Discussion

We investigated genome-wide autozygosity patterns and genomic inbreeding levels based on homozygosity-by-descent segments and estimate the genetic parameters for a self-fertilized population of *Eucalyptus*. In addition, we calculate inbreeding depression at the family level considering a mixed matting system model and used simulations that emulated our data to evaluate the relative bias of inbreeding depression estimators due to unbalance among self-fertilized and cross-fertilized individuals in the families.

### 4.1. Genetic Parameters

The inclusion of dominance genetic effects in genomic prediction models enabled a more precise partitioning of genetic variance. When dominance effects are not considered, additive effects are often overestimated (Dias et al., 2018). By incorporating dominance into our models, we achieved improved predictive accuracy and a more realistic representation of the genetic contributions underlying the expression of DBH. Studies have demonstrated that dominance variance largely contribute to the phenotypic variation of growth traits in *Eucalyptus* (Martins et al., 2024). For instance, the additive- dominant genomic model improved predictions for growth (mean annual increment) by 5 to 14% compared to an additive genomic model, although no improvement was seen for wood quality (Resende et al., 2017a).

In Eucalyptus hybrids, dominance has been identified as a significant contributor to growth traits, with dominance-to-additive variance ratios exceeding 1.2 in certain hybrids (Tan and Ingvarsson, 2022). Incorporating dominance effects into genomic selection models has been shown to improve predictive accuracy, underscoring the importance of considering non-additive genetic effects in tree breeding strategies (Thavamanikumar et al., 2020; Muñoz et al., 2014). In the presence of directional dominace, a phenomenon where the average effect of dominance deviations across loci tends to favor the expression of the heterozigote, heterosis can occur (Wright, 1984). Heterosis is when the offspring of two genetically distinct inbred lines exhibit superior traits compared to their parents. Although we performed only one generation of selfing, we can assume that in some level the principles defined by Shull (1908, 1910) and East (1909) for maize hold to our population. Shull’s works demonstrated that selfing, or inbreeding, could isolate homozygous lines, which, when crossed, could capitalize on heterosis (Shull, 1908, 1910). This hypothesis was supported by East, who applied Mendelian genetics to the study of heterosis. East suggested that homozygosity in inbreds enhanced deleterious effects, which were mitigated when inbreds were hybridized (East, 1909). In addition to dominance effects, heterosis can also be driven by complementarity, arising from additive gene actions (Retief and Stanger, 2009; Potts and Dungey, 2004). Therefore, the performance of hybrids originated from the latter generations of selfing, may validate the hypothesis that this same behavior associating inbreeding and heterosis occurs in eucalyptus.

### 4.2. Estimation of inbreeding coefficient

In a single generation of self-fertilization (without selection), assuming that the parent is heterozygous, it is theoretically expected that 50% of the loci in the progeny would remain heterozygous while the other 50% would become homozygous, resulting in an inbreeding coefficient of 0.5. Contrary to this expectation, in our population, we found an estimated inbreeding coefficient of 0.2367, suggesting strong selection against homozygous individuals. Myburg et al. (2014) observed an inbreeding coefficient of 0.355 in progenies derived from self-fertilized parental genotypes or *Eucalyptus grandis*. Different patterns of homozygosity and heterozygosity have been reported across various plant species Somera et al. (2018). In natural trees, inbreeding coefficients were found to decrease with age, from 0.3 in young plants to 0.15 in adults ones, suggesting that natural selection may reduce the frequency of inbred individuals over time (Angbonda et al., 2024).

### 4.3. Inbreeding depression

Inbreeding depression from self-fertilization was studied in *Eucalyptus globulus* (Costa e Silva et al., 2010) using self-fertilized, open-pollinated, and hybridized plants with data from 4, 6, and 10 years. Costa e Silva et al. (2010) found that self-fertilized trees exhibited lower survival rates than open-pollinated ones and the survival difference between self-fertilized and open-pollinated plants increased over time. By the age of ten years, survival rates were 21.9% for self-fertilized while 82.9% for open-pollinated trees. Bison et al. (2006) made a comparison between open-pollinated progenies and hybrids performance in *E. grandis* and *E. urophylla* and found a mean inbreeding depression for circumference at breast height equals to 17.5% for *E. grandis × E. urophylla*, which is smaller than found for our data, since they were not working with self-fertilized populations. In both studies, the early evaluation of these trees can contribute to an overestimation of the *ID*, since it tends to increase as age advances. Costa e Silva et al. (2011) examined different levels of inbreeding in *E. globulus* and observed both strong age and environmental effects on inbreeding depression. Inbreeding depression was also observed for oil palm (Luyindula et al., 2005), macaw palms (Lanes et al., 2015), and Cocos nucifera (Pandin, 2009) trees decreasing the mean of vegetative and productive traits.

Inbreeding depression tends to be greater for growth than for seed traits, especially for those from self-fertilized individuals (Pereira et al., 2020; Aguiar et al., 2020), which may be due to the purging of inbred seeds, which did not germinate due to postzygotic mechanisms. Since we selected 30 seedlings from each of the 20 female parents after the germination phase, the early effects of inbreeding may have been masked in our population. In addition, there may have been a selection of healthier seedlings, which may have increased the representation of less inbred individuals. On top of this, harmful and deleterious alleles are expected to be rare in tree populations (Namkoong and Bishir, 1987).

Inbreeding depression can alter the expression of growth and survival traits, affect genetic parameters, and reduce selection responses, with its impact intensifying over time (Costa e Silva et al., 2010). Our findings highlight the need to maintain genetic diversity in breeding populations to mitigate inbreeding depression. Strategies such as reciprocal recurrent selection can enhance genetic divergence between populations, increasing genetic variance (reducing the negative impact of inbreeding depression), and exploiting heterosis (Chaves et al., 2024).

### 4.4. Simulation

Our results suggest that the inbreeding depression estimator is particularly sensitive to the unbalance in the composition of families. As the proportion of selfed individuals increases, the bias becomes more pronounced, particularly when fewer than seven crossed individuals are included in the family composition. One of the primary challenges posed by unbalanced sample sizes is the potential distortion of statistical power and effect size estimates. Studies have shown that the performance of statistical methods can degrade as the degree of sample imbalance increases (Yang et al., 2006). This degradation can lead to biased results, particularly in hypothesis testing, where the power to detect true effects may be compromised.

Moreover, the variance among different statistical methods becomes more pronounced with increased sample imbalance (Yang et al., 2006). This variance suggests that certain methods may be more robust against imbalance, highlighting the importance of method selection in research design.

In more balanced configurations (e.g., 23.7 through 15.15), both variability and *RMSE* values decline significantly. Therefore, our findings reveal that configurations with a pronounced imbalance between self and cross- fertilized individuals exhibit heightened variability and *RSME* values, indicating reduced estimation accuracy. This observation is related to previous research that demonstrates that small sample sizes can lead to unstable mean estimates, thereby amplifying errors in *ID* calculations (Pérez-Pereira et al., 2023). In fact, the performance of statistical methods is expected to degrade as the degree of sample imbalance increases. This degradation can lead to biased results, particularly in hypothesis testing, where the ability to detect true effects may be compromised (Kun Yang, 2006). Therefore, our results offer practical guidance for experimental planning, where moderate imbalances may suffice to produce robust estimates of inbreeding depression, reducing logistical challenges without compromising data reliability.

There are some limitations of ours study that deserve to be mentioned. First, due to the large cycle of Eucalyptus, there is only one generation of selfing available, which prevents us from making reliable estimates of the inbreeding in more advanced generations of selfing. Furthermore, there is a reduced number of individuals per family, and the number of selfed and crossed individuals per family were uneven, which could impact the accuracies. In addition, the trial was established only in one environment, which can make broader interpretations difficult. Finally, there is room for refinements in the data simulation process since after testing several scenarios and cycles and a large number of repetition cycles we achieved an overlay of 89%. Our study has certain limitations that warrant discussion. First, due to the large cycle of Eucalyptus, there is only one generation of selfing available, which prevents us from making reliable estimates of the inbreeding in more advanced generations. Additionally, the number of individuals per family was relatively small, and there was an imbalance in the number of selfed and crossed individuals per family, which may have affected the accuracy of our estimates. Moreover, the trial was conducted in a single environment, which may limit the interpretation of our findings for other evironmental conditions. Lastly, while our data simulation process included multiple scenarios, cycles, and a large number of repetitions, there is still room for refinement. Despite these efforts, we achieved an overlap of 89% among real and simulated data, indicating potential areas for improvement in the simulation methodology.

## 5. Conclusion

Our study presents a comprehensive investigation into the genetic consequences of self-pollination in *Eucalyptus* spp. providing a detailed investigation into genetic parameter estimation, inbreeding depression, and biases from unbalanced groups. We found significant contributions of both additive and dominance variance to diameter at breast height, with broad-sense heritability at 0.38 and narrow-sense heritability at 0.24, emphasizing the importance of genetic components in trait expression. Inbreeding depression, evident in most families, was associated with reduced DBH as inbreeding levels (*F*_*HBD*_) increased. Families with lower inbreeding exhibited higher genotypic values for DBH, highlighting the need to manage homozygosity to maintain productivity when family selection is performed. Simulations showed that unbalanced sample sizes of selfed and crossed individuals over-estimate inbreeding depression, with accuracy improving as sample balance increased. By knowing these genomic patterns, breeders can strategically combine selfing and hybridization to explore genetic variability, mitigate inbreeding depression and accelerating genetic gains. Future research should focus on extending these analyses to later generations of inbreeding, incorporating interactions with environmental factors and multiple environments, and exploring the role of epistasis in trait expression. Such efforts will ensure the viability of this breeding strategy in eucalyptus and other perennial species.

## 6. Author contribution statement

Ferreira, F.M., Tambarussi, E.V., and Dias K.O.D., designed the research. Ferreira, F.M. and Netto J.A.F.V performed the statistical analysis. Ferreira, F.M., Netto J.A.F.V, and Melchert G.F. wrote the first draft. Tambarussi, E.V., Dias K.O.D., Bhering L.L., Muniz, F.R, Benatti, T.R, and Matos, J.W. revised the drafts.Matos, J.W., Muniz, F.R, and Benatti, T.R, data curation. All the authors read and approved the final manuscript.

## 7. Acknowledgements

This study was financed, in part, by the São Paulo Research Foundation (FAPESP), Brazil. Process Number #2023/04881-3. Tambarussi E.V. was supported by CNPq (research productivity fellowship 407175/2021-0). The authors would like to thank the company Suzano S.A. for providing the data for this work.

