## Supplementary Material for "Unraveling the inbreeding depression patterns in a self-pollinated eucalyptus population"

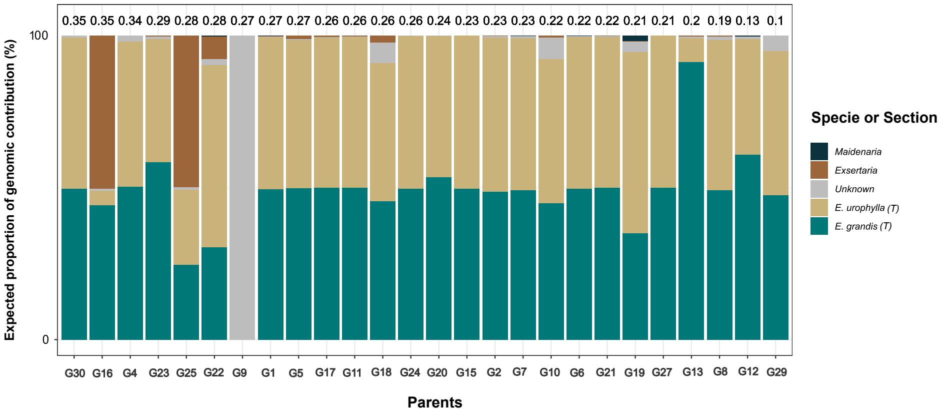


Figure S1: The expected proportion, in percentage, of each species or section in the composition of the male parent via admixture analyses. The letter T represents the section Transversaria. The values above each bar represent the inbreeding coefficients calculated via the homozygosity-by-descent approach ($F_{HBD}$). The parents are arranged from highest to lowest $F_{HBD}$.

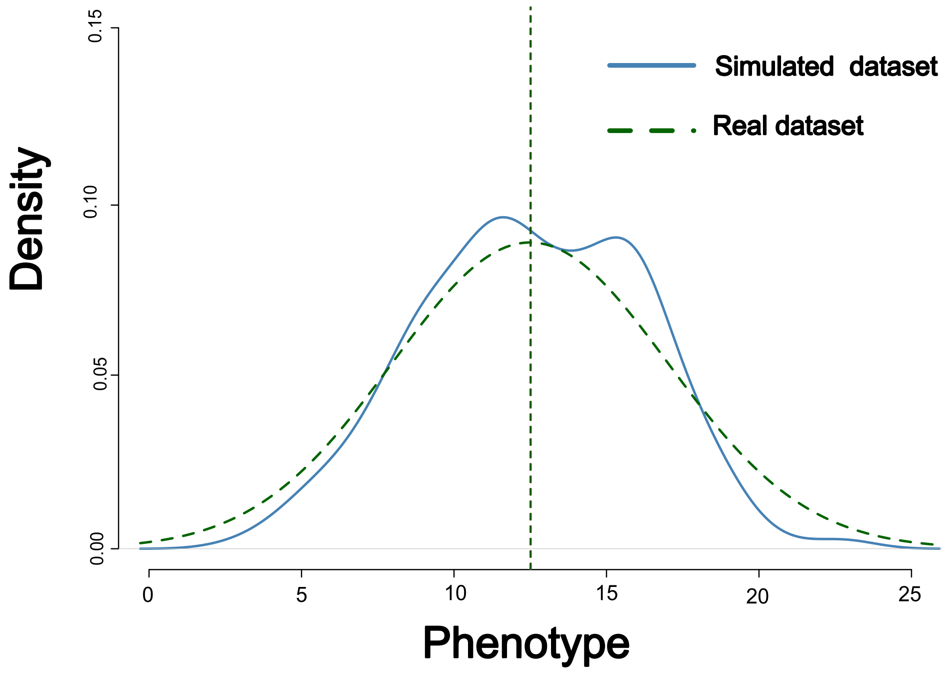


Figure S2: Overlay between simulated and rela dataset of a $S_{0:1}$ population of *Eucalyptus* spp.

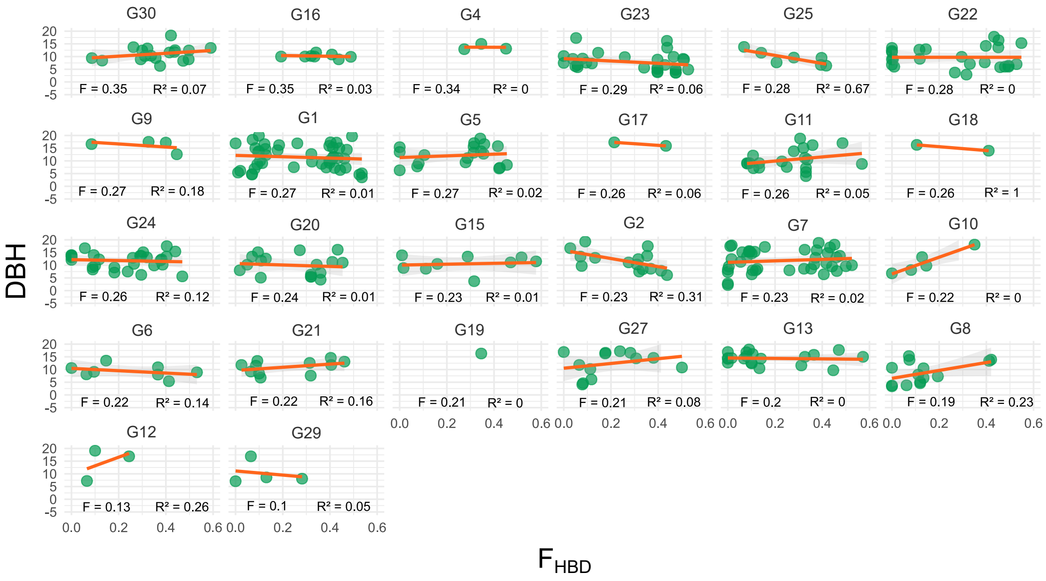


Figure 3: Regression of Diameter at Breast Height (DBH) on $F_{DBH}$ for each evaluated family.
